# A semi-automated cell tracking protocol for quantitative analyses of neutrophil swarming to sterile and *S. aureus* contaminated bone implants in a mouse femur model

**DOI:** 10.1101/2023.12.07.570663

**Authors:** Sashank Lekkala, Youliang Ren, Jason Weeks, Kevin Lee, Allie Jia Hui Tay, Bei Liu, Thomas Xue, Joshua Rainbolt, Chao Xie, Edward M. Schwarz, Shu-Chi A. Yeh

## Abstract

Implant-associated osteomyelitis remains a major orthopaedic problem. As neutrophil swarming to the surgical site is a critical host response to prevent infection, visualization and quantification of this dynamic behavior at the native microenvironment of infection will elucidate previously unrecognized mechanisms central to understanding the host response. We recently developed longitudinal intravital imaging of the bone marrow (LIMB) to visualize fluorescent *S. aureus* on a contaminated transfemoral implant and host cells in live mice, which allows for direct visualization of bacteria colonization of the implant and host cellular responses using two-photon laser scanning microscopy. To the end of rigorous and reproducible quantitative outcomes of neutrophil swarming kinetics in this model, we developed a protocol for robust segmentation, tracking, and quantifications of neutrophil dynamics adapted from Trainable Weka Segmentation and TrackMate, two readily available Fiji/ImageJ plugins. In this work, *Catchup* mice with tdTomato expressing neutrophils received a transfemoral pin with or without ECFP-expressing USA300 methicillin-resistant *Staphylococcus aureus* (MRSA) to obtain 30-minute LIMB videos at 2-, 4-, and 6-hours post-implantation. The developed semi-automated neutrophil tracking protocol was executed independently by two users to quantify the distance, displacement, speed, velocity, and directionality of the target cells. The results revealed high inter-reader reliability for all outcomes (ICC > 0.98; p > 0.05). Consistent with the established paradigm on increased neutrophil swarming during active infection, the results also demonstrated increased neutrophil speed and velocity at all measured time points, and increased displacement at later time points (6 hours) in infected versus uninfected mice (p < 0.05). Neutrophils and bacteria also exhibit directionality during migration in the infected mice. The semi-automated cell tracking protocol provides a streamlined approach to robustly identify and track individual cells across diverse experimental settings and eliminates inter-observer variability.

## Introduction

Implant-associated osteomyelitis remains one of the most prevalent and serious orthopaedic problems: the incidences of infection for all orthopaedic subspecialties range from 0.1%-30%, at a cost of $17,000-$150,000 per patient.(1) As neutrophil swarming to the surgical site is a critical host response to prevent infection, visualization and quantifying this dynamic behavior at the native microenvironment of infection will provide previously unrecognized mechanisms central to understanding the host response that may not be fully recapitulated ex vivo.(2)

Advances in intravital microscopy of long bones, such as the utilization of Gradient Index (GRIN) lenses, have facilitated deep-tissue visualization including longitudinal imaging of the bone marrow (LIMB) deep within the femur.(3) Specifically, we previously described LIMB which allows direct visualization of fluorescent host cells and bacteria proximal to a transfemoral implant in mice.(4) These imaging systems facilitate the investigation of immune mechanisms, pathogen evasion strategies, and the effects of different therapeutic agents on the kinetics of immune cells. To quantify neutrophil swarming kinetics, TrackMate(5,6), an open-source, user-friendly plugin available within Fiji(7) is an attractive option that allows interactive cell tracking. Previous studies have reported diverse approaches to counting and tracking cells using TrackMate.(8–12) However, some of these tracking approaches utilize computationally expensive models or are restricted to specific operating systems.(12) The most apparent cellular architecture in the bone infection setting is the accumulation of bacteria and host immune cells over time. In this regard, these protocols are sub-optimal in segmenting and tracking densely populated cells with variable shapes and fluorescence intensities which became even more problematic in the context of three-dimensional intravital multiphoton laser scanning microscopy (IV-MLSM). While manual tracking options are available,(6,12) they are subject to significant inter-observer variation. Subsequent statistical analyses are therefore not meaningful given such inter-user variability.

To overcome these limitations, we developed post-processing workflow, and protocols taking advantage of Trainable Weka Segmentation (TWS)(13), a machine learning tool available within Fiji, to accurately segment the cells from the background before feeding the data to TrackMate. This automatic segmentation eliminated the need for user-defined variables, such as thresholding, noise removal, etc., and largely minimized inter-user variability. Moreover, a classifier trained by the user to recognize the cells of interest can be created using TWS, which can be used to batch-process videos for high-throughput analyses. The developed protocol enabled robust cell tracking in IV-MLSM timelapse videos of neutrophils adjacent to infected and uninfected femoral implants in vivo, and their interactions with bacteria.

## Methods

### MRSA strain and implants

The most prevalent community-acquired methicillin-resistant *Staphylococcus aureus* (MRSA) strain, USA300, was used for all experiments. We transformed USA300 LAC (ATCC AH1680) with pCM29-*sarA*::*ecfp* reporter plasmid and pCM29-*sarA*::*egfp* reporter plasmid to generate ECFP and EGFP expressing USA300, respectively. The transformed bacteria were positively selected using chloramphenicol (10μg/ml).(4)

### Animal Surgery and LIMB

All animal research was performed under protocols approved by the University of Rochester Committee on Animal Resources (UCAR-2019-015). The surgical procedure and the imaging setup were based on protocols previously described.(4) Briefly, the right femur was implanted with a customized LIMB system. The implant enabled imaging of a fixed region of interest (ROI) proximal to an L-shaped transcortical pin (3mm long) in the diaphyseal bone marrow. Catchup mice(14) with tdTomato expressing neutrophils received either a sterile pin or a pin incubated in overnight cultures of ECFP^+^ MRSA for 30 minutes prior to the implantation procedure.

### Intravital two-photon laser scanning microscopy

IV-MLSM was performed using an Olympus FVMPE-RS multiphoton imaging system (Olympus), equipped with MaiTai and InsightX3 Titanium:Sapphire lasers (Spectra-Physics, Santa Clara, CA), and an LUCPLFLN 20× (NA 0.45) air objective (Olympus, Tokyo, Japan). The lasers were tuned to 1050 nm and 860 nm and the fluorescence of tdTomato and ECFP were collected with 575−630 nm and 460−500 nm filters, respectively. Images were acquired at a size of 512 × 512 pixels with a 0.01-μs pixel dwell time. At 2, 4, and 6 hours after implantation, three-dimensional (424.26×424.26×60 μm^3^), time-lapse (30-second interval for 30 minutes) image stacks were acquired using FLUOVIEW (31S-SW, Olympus).

### Cell tracking and image quantification

We developed a semi-automated protocol to track neutrophils and quantify their characteristics of cell motility (Fig 1A, see Supplementary Methods for a step-by-step guide). In brief, maximum intensity projection was used to project the three-dimensional image stacks in 2D given that the cells of interest and bacteria are primarily on the surface of the implant. The image stacks were drift-corrected using the Image Stabilizer plugin.(15) To remove noise and non-moving artifacts contributed by autofluorescent objects, the minimum intensity projection of the time-lapse images was subtracted from all the slices (Fig 1C). To avoid inter-user variability primarily introduced by inaccurate cell identification during the tracking process, the neutrophils were first segmented using TWS, which uses a library of machine learning training features to generate a probability score of the foreground (Fig 1D).(13) The training features for the TWS classifier can be categorized into noise removal, edge detection, and texture description features.(13) We compared various combinations of these features focusing on noise removal and edge detection to optimize cell detection and tracking (S2 Appendix). The best-performing classifier incorporated Gaussian blur, Sobel filter, Hessian, and Difference of Gaussians training features with a minimum sigma of 1 and a maximum sigma of 16. For classification, we used Fast Random Forest, the default machine learning model in TWS. After identifying the best training features, we refined the labels for background and neutrophils to arrive at an optimal classifier which was consistently employed for all timelapse videos. The probability maps generated were then used as input for TrackMate to track moving cells.

**Fig 1.**
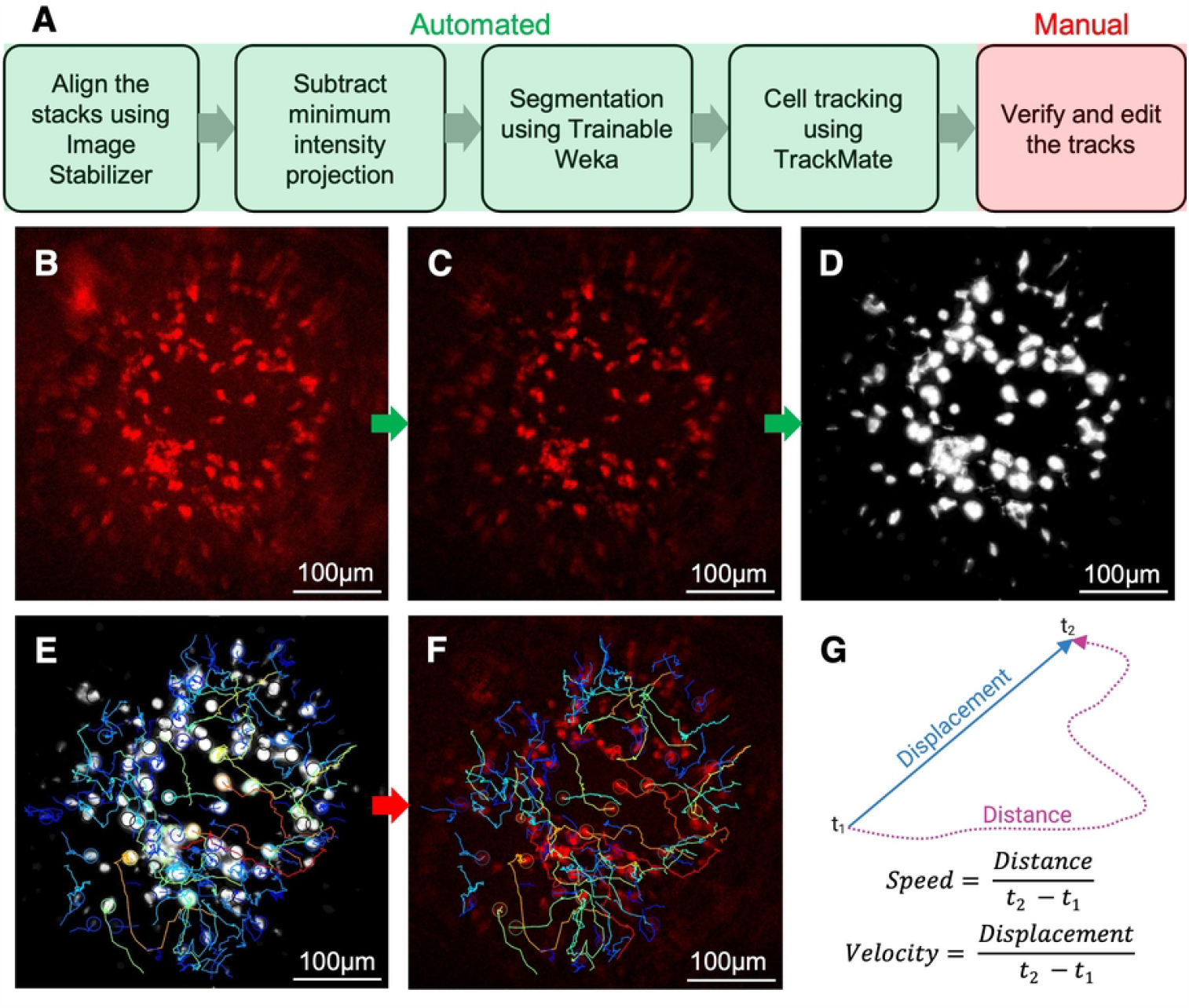
Workflow for cell tracking using trainable Weka segmentation and TrackMate in ImageJ. (A) Flowchart of the protocol. (B) Representative 2D LIMB image from a 30-minute timelapse video (Scale bar = 100μm) (C) The minimum intensity projection of the timelapse stack was subtracted from all the slices to reduce noise and remove non-moving artifacts. (D) A custom classifier trained to detect neutrophils was applied to create probability maps using Trainable Weka Segmentation (TWS). (E) In TrackMate, the cells were detected using a Laplacian of Gaussian (LoG) detector, and the tracks were determined by a linear assignment problem-based (LAP) tracker. (F) These tracks were overlaid on (C) and edited manually for accuracy. (G) A custom MATLAB code was used to calculate the distance and displacement for each neutrophil track. Speed and velocity were calculated as the ratios of distance and displacement to track duration, respectively. Directionality was calculated as the ratio of displacement and distance. Representative images from N=6 mice.

Specifically, the moving objects are defined in TrackMate based on cell diameter and total displacement. The cells were then detected using a Laplacian of Gaussian (LoG) detector, and the tracks were determined by a linear assignment problem-based (LAP) tracker (Fig 1E).(5,6). The maximum linking distance was set to 20μm based on ground truth provided by manual tracking. Note that this distance is determined empirically by the time interval of longitudinal imaging (30 seconds in our case). After the tracks were computed, a displacement filter of 16μm was set to remove tracks that are a result of minor intensity fluctuations in stationary cells. These tracks were overlaid on the drift-corrected stack and edited manually as needed for accuracy (Fig 1F). A custom MATLAB code (S3 Appendix) was used to calculate the distance and displacement for each neutrophil track. Speed and velocity were calculated as the ratios of distance and displacement to track duration, respectively (Fig 1G). Directionality, a measure of the straightness of a track was calculated as the ratio of displacement and distance.

Using this protocol, we also tracked bacteria by employing a different TWS classifier and changing the particle size to 8μm. To study the migration pattern and interactions of neutrophils and bacteria, a custom MATLAB script was developed to map and track spatial associations of neutrophils and bacteria based on their migration trajectories (S4 Appendix). The script also identifies cell interaction events based on criteria that the neutrophils and bacteria were in proximity (within 16μm distance) for a prolonged duration (≥ 5 time frames, 150 seconds).

The volume of neutrophils proximal to the implant was quantified using Imaris (Oxford Instruments). Briefly, the 3D stack was smoothed, the background was subtracted, and a user-defined threshold was applied to segment neutrophils. Then the imaging artifacts were manually removed, and the neutrophil volume was calculated based on the number of voxels (S1 Fig, Supplementary Methods).

### Statistics

Data are presented as median ± interquartile range. Mann-Whitney tests adjusted for multiple comparisons by the Holm-Šídák method were used to test if the tracking metrics differed between individuals and the study groups. Interclass correlation coefficients (ICC) were calculated based on two-way random effects model with absolute agreement for two raters.(16)

## Results

### Reproducibility of the tracking protocol

The major limitation of manual tracking is high inter-user variability. The two users assigned to analyze the same IV-MLSM video often identified different tracks. The lack of consistencies is shown in the data distribution, where the median displacement and velocity of neutrophils differed by 25% and 22% respectively between the two users (S2 Fig). While these results were not statistically significant, such substantial numerical differences impede reliable conclusions.

In contrast, our protocol yields reproducible tracking metrics from two users who independently analyzed three IV-MLSM videos. While there were minor differences in the tracks generated by both users (Fig 2A, 2B), we obtained robust inter-reader reliability (ICC > 0.98). The track metrics and data distribution are in good agreement (p > 0.05) (Fig 2C-2G).

**Fig 2.**
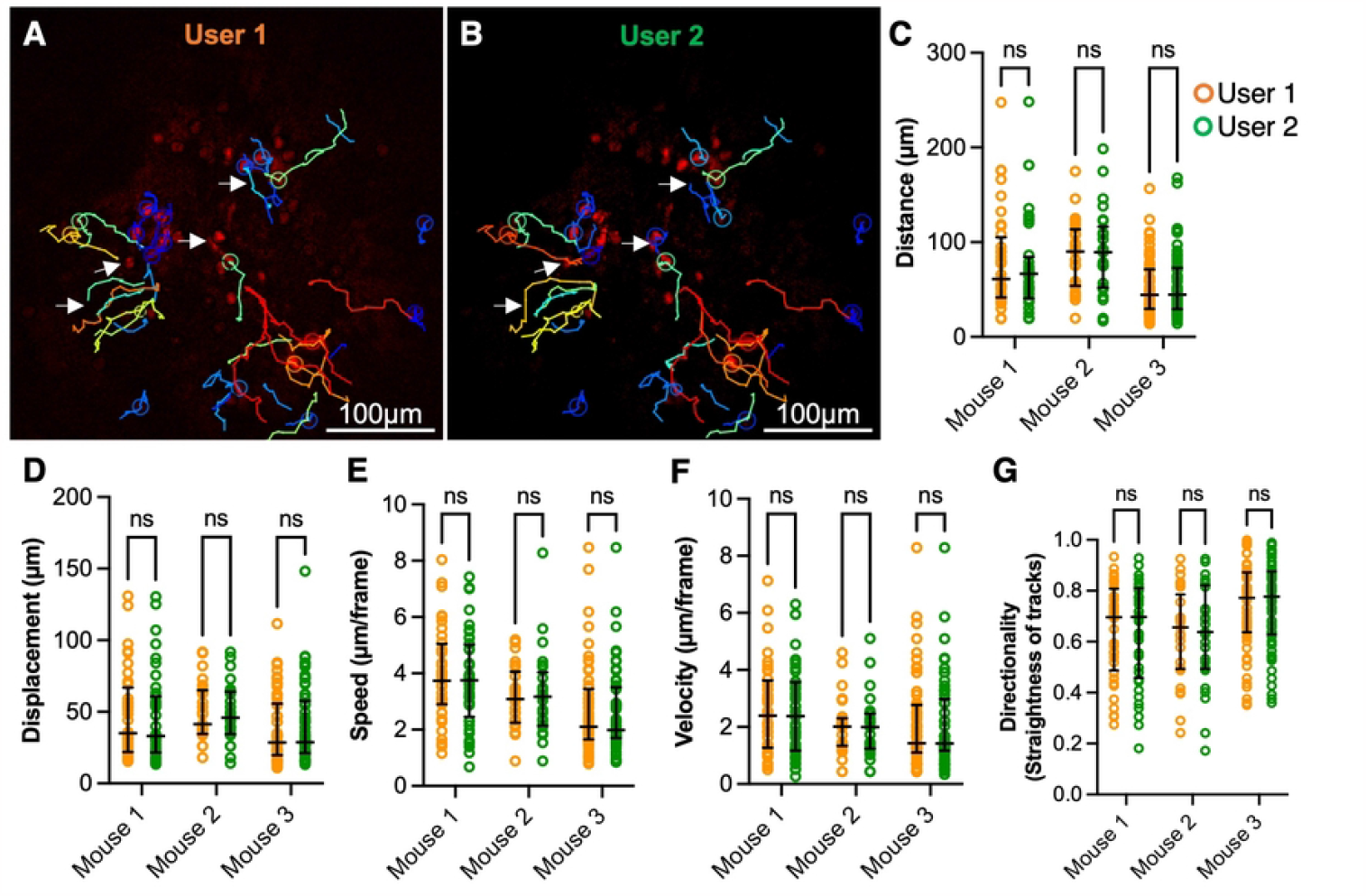
The tracking protocol resulted in low inter-person variability in generating tracks. Two users independently analyzed three IV-MLSM timelapse videos and the generated track parameters were compared. Representative 2D IV-MLSM image with overlaid tracks generated by user 1 (A) and user 2 (B). The differences in the tracks generated by both users are shown with white arrows. Semiautomated quantification of neutrophil distance traveled (C), displacement (D), mean speed (E), mean velocity (F), and directionality (G), was performed and the data are presented with the median and interquartile range. (**\***p < 0.05 as determined by Mann-Whitney tests adjusted by the Holm-Šídák method (n=25-57 tracks. Representative images from N=3 mice)).

### Quantification of neutrophil swarming proximal to infected vs. sterile implant

As it has been well established that neutrophils demonstrate increased swarming in the setting of an active infection,(2,17,18) we validated our protocols by comparing neutrophil behavior proximal to MRSA infected versus sterile femoral implants. Consistent with the infection status, the neutrophil volume proximal to the pin increased with time in the infected animals but remained consistent in uninfected animals (Fig 3A-B, S1-3 Videos). The neutrophils in the infected animals traveled longer distances at 6 hours and longer displacements at 4 hours and 6 hours compared to the neutrophils in uninfected animals (Fig 3C-D). Despite the path length of migration increasing several hours after implantation, the neutrophils in the infected animals traveled at significantly greater speeds (distance/time) and velocities (displacement/time) at all time points compared to uninfected animals (Fig 4E-4F). In agreement, the directionality of the neutrophil migration in the infected animals, indicative of swarming behaviors, was shown at the early time point (2 hours) compared to uninfected animals (Fig 4G).

**Figure 3.**
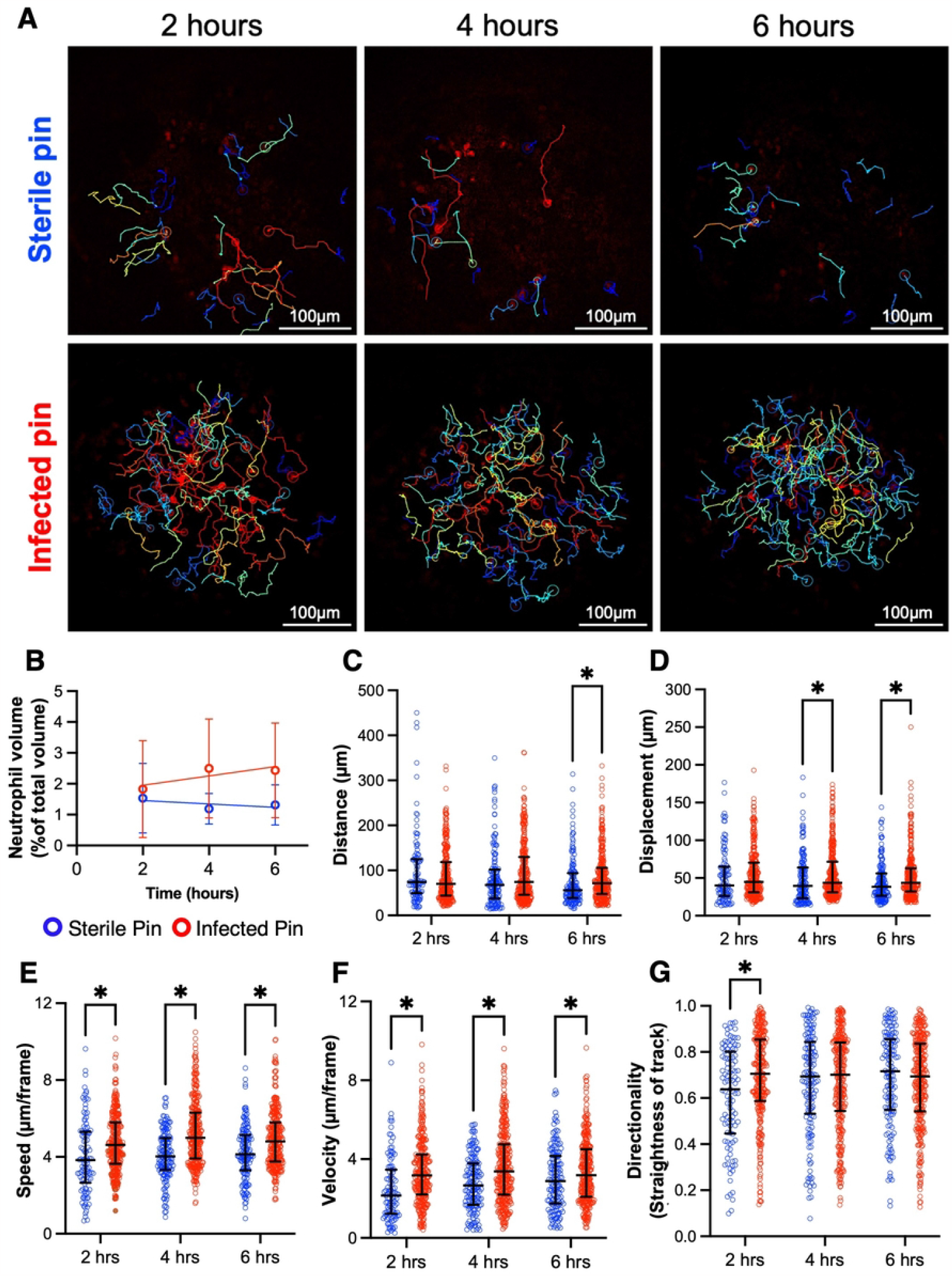
Increased neutrophil swarming proximal to MRSA contaminated vs. sterile bone implant. *Catchup* mice were challenged with a sterile or MRSA-contaminated transfemoral implant and neutrophil swarming behaviors were quantified from 30min IV-MLSM videos obtained at the indicated time post-implantation. (A) Representative 2D IV-MLSM images with overlaid neutrophil tracks proximal to sterile and infected pin at 2-, 4-, and 6-hours post-implantation (S1-3 Videos). (B) Change in neutrophil volume with time proximal to infected and sterile implants calculated using Imaris (S1 Fig). Semiautomated quantification of neutrophil distance traveled (C), displacement (D), mean speed (E), mean velocity (F), and directionality (G), was performed and the data are presented with the median and interquartile range. (*p < 0.05 as determined by Mann-Whitney tests adjusted by the Holm-Šídák method (n=105-316 tracks/group/timepoint, N=3 mice/group)).

**Fig 4.**
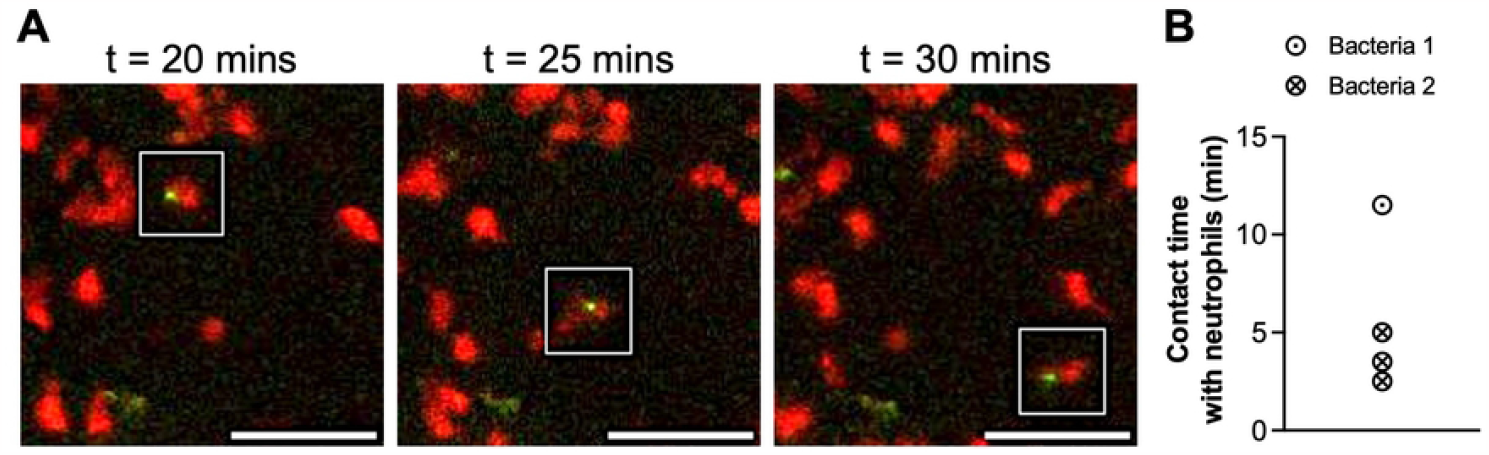
Parallel migration pattern of neutrophils and bacteria. A custom script quantifies the interactions of neutrophil and bacteria tracks. (A) Representative 2D IV-MLSM images showing parallel migration of neutrophils and bacteria over several time points where the cells made stable contact with the neutrophil (Scale bar = 50μm). (B) Contact time between neutrophils and bacteria revealed stable interaction or intermittent contact with several neutrophils (S4-5 Videos).

Moreover, by tracking both bacteria and neutrophils, parallel migration pattern was observed (Fig. 4A), where bacteria may exhibit stable contact with a neutrophil, or made intermittent contact with multiple cells (Fig 4B). The contact time and frequency may be a potential metric at the early time point to correlate with phagocytosis events at later time points (colocalization of neutrophils and bacteria) and the efficacy of different therapeutic regimens on neutrophil-pathogen interactions.

## Discussion

In this paper, we described a protocol to track cells from IV-MLSM videos accurately and reproducibly. The protocol included post-processing steps to minimize stationary and motion artifacts and employed TWS, a machine learning algorithm for robust segmentation of the target cell population. The use of these segmented images greatly reduced the inter-user variability when generating tracks in TrackMate and customized Matlab code. Using these quantitative metrics, we showed that neutrophil swarming is more pronounced proximal to infected implants compared to uninfected implants.

This protocol is semi-automated with minimal user interference which resulted in high reproducibility (Fig 2). By standardizing the pre-processing and segmentation methods, the need for selecting an arbitrary threshold, a main source of inconsistencies, was eliminated. Further, utilizing a singular global threshold might not yield optimal results for neutrophil segmentation due to the inherent variability and signal to noise ratios in fluorescence associated with 3D intravital imaging. TWS overcomes this limitation by using a collection of machine learning algorithms to produce pixel-based segmentation. In addition to automating the process, we have outlined a set of guidelines for manual track verification (Supplementary Methods) which further reduces inter-user variability.

We opted not to extensively denoise the image prior to segmentation to retain as many morphological and textural features as possible. Such information is a critical “training set” to identify the background in the machine learning algorithms. We subtracted the minimum intensity projection from the stack to remove any stationary artifacts. However, this method will also remove any stationary cells and the nidus. Given our focus on tracking only the moving cells, this pre-processing method proved most effective for our purpose. This protocol can be adapted with alternative denoising techniques such as morphological filtering (for example, white top-hat) to preserve the stationary details.(19) Depending on the type of noise and artifacts, images can be directly segmented using TWS without any pre-processing.

Note that we chose a conservative classifier in TWS for its capability to detect even faintly fluorescent neutrophils. Consequently, the resulting probability maps depicted larger neutrophils compared to the original stack (Fig 1C, 1D). The approach is sufficient in detecting early swarming behaviors after the onset of infection. A potential drawback of this classifier is that when the cells are densely packed, potentially at even later time points, multiple neutrophils may appear as a single object. A more aggressive classifier may perform better in distinguishing individual cells although a trade-off of missing dim subjects is expected (S3 Fig). Deep learning programs such as StarDist which also use object boundaries for segmentation may overcome this limitation with a custom training set.(10,20) However, StarDist is most effective for cases of nuclear staining or cells with star-convex shapes which may not always apply to varying shapes of patrolling cells such as neutrophils.(20)

The probability maps were used to generate tracks in TrackMate. For object detection, a Laplacian of Gaussian (LoG) detector which detects local maxima was used.(21) Note that the object size was set to 16μm, slightly larger than typical sizes of neutrophils (< 8-15 μm). This is based on size fitting on several probability maps to ensure separate detection of two closely spaced neutrophils. A quality filter (= 0.001) which is a measure of the value of the local maxima and the size of the object was applied to remove objects detected from noise. For linking these objects, we employed a linear assignment problem-based algorithm (LAP), which links objects based on the lowest square distance.(22) Note that this approach results in an expected limitation when two cells come close to each other, the algorithm may favor linking these two cells at the frame of contact instead of tracking the same cell. These discrepancies were manually corrected.

Speed and velocity are the most reliable indicators of cell kinematics, as distance and displacement can be influenced by the number of frames a cell was tracked, cell density, and potential travel beyond the imaging session’s duration. Additionally, it is important to note that shorter tracks tend to be straighter, which potentially biases the distribution of the directionality parameter.

Overall, an inherent limitation of all cell tracking protocols is that they perform less optimally when the density of cells is high.(23) Our protocol faces the same limitation and some of the cells were not tracked by the protocol (S1-4 Videos). In this study, user verifications of error-prone regions (e.g., cells with small displacement, low intensity, or located at the boundaries of the field of view) are necessary. Each track took about a minute for the users to verify (across 60 frames). Previous studies proposed solutions to overcome this limitation by introducing specific rules for linking tracks when two cells collide or come close to each other.(8) However, these algorithms are computationally expensive, and these rules are specific to the cell type and imaging settings. Future optimization focusing on connection rules and penalties will help address this issue.

Altogether, we developed a semi-automated tracking protocol with minimum user interference by leveraging the capabilities of TWS and TrackMate, two readily available Fiji plugins. We showed that the protocol offers low inter-user variability along with high accuracy. The protocol can be easily adapted to study the kinematics of different cells by changing the TWS classifier and the tracking parameters. Overall, our cell tracking protocol offers a streamlined approach to tracking individual cells accurately and efficiently in diverse experimental settings.

## Acknowledgments

Schematic images were generated with BioRender. The authors thank Kaye Thomas, Emma Norris, Julie Zhang, Yurong Gao, and Rhonda Jean Kay for their assistance with the LIMB imaging and analyses.

## Supplementary Information

### Supplementary Methods

**Step-by-Step Protocols for Cell tracking using Trainable Weka Segmentation and TrackMate**

**Volume analysis using Imaris**

## Supplementary Figures

**Supplementary Figure 1. Workflow for volumetric quantification of neutrophil volume using Imaris**.

**Supplementary Figure 2. Inter-person variability before using the semi-automated protocol**.

**Supplementary Figure 3. Comparison of two different TWS classifiers**.

**Supplementary Video 1**. Representative IV-MLSM timelapse of the neutrophils with the tracks overlaid proximal to a sterile pin (left) and an infected pin (right) at 2 hours. Scale bar = 100μm. Note that the undetected neutrophils have displacement lower than 16μm.

**Supplementary Video 2**. Representative IV-MLSM timelapse of the neutrophils with the tracks overlaid proximal to a sterile pin (left) and an infected pin (right) at 4 hours. Scale bar = 100μm. Note that the undetected neutrophils have displacement lower than 16μm.

**Supplementary Video 3**. Representative IV-MLSM timelapse of the neutrophils with the tracks overlaid proximal to a sterile pin (left) and an infected pin (right) at 6 hours. Scale bar = 100μm. Note that the undetected neutrophils have displacement lower than 16μm.

**Supplementary Video 4**. Representative IV-MLSM timelapse of the neutrophils (left) and USA300 MRSA (right) with the tracks overlaid proximal to an infected pin at 2 hours. Scale bar = 100μm. Note that the undetected neutrophils and bacteria have displacement lower than 16μm.

**Supplementary Video 5**. Representative IV-MLSM timelapse of the neutrophils and bacteria proximal to an infected pin at 2 hours (same experiment as Supplementary Videos 1 and 2). The insets are tracking two different bacteria. Scale bar = 100μm. Note that the undetected neutrophils and bacteria have displacement lower than 16μm.

**S1 Appendix – ImageJ macro for pre-processing**

**S2 Appendix – TWS classifier**

**S3 Appendix – MATLAB code for quantification of tracks**

**S4 Appendix – MATLAB code for quantification of parallel migration pattern**

